# Dual-Logistic Analysis of Time- and Concentration-Dependent Phenotypic Efficacy Evaluation Integrating Drug Targets Information

**DOI:** 10.64898/2026.03.09.709547

**Authors:** Lingli Wang, Rumeng Qu, Qialing Huang, Min Hu, Tongsheng Chen

## Abstract

Tumor heterogeneity highlights the necessity of precision cancer medicine, making the evaluation and screening of anticancer drugs a core challenge in cancer therapy. However, current cell-based efficacy assessment methods struggle to quantify the holistic impact of drugs on cellular behavior through specific target engagement. Here, we proposed a novel approach (DL-TCP-FRET) that integrates phenotypic and target-related evaluations: the logistic fitting analysis is performed on time- and concentration-dependent cellular phenotypic characteristics to construct a phenotypic score (*P*), while a target score (*T*) is established based on the FRET efficiency between target proteins. These two scores were then further combined to generate a unified drug efficacy score (*PT)*. Validation in A549 cells demonstrated that our method can reliably distinguish EGFR-TKIs from non-targeted drugs. DL-TCP-FRET simplifies the experimental workflow of drug efficacy evaluation and improves the accuracy of targeted drug identification, providing a novel strategy for advancing precision cancer therapy.

## 1. Introduction

Precision clinical medication has emerged as a pivotal strategy in current cancer therapy, facilitating the transition of therapeutic approaches from empirical to personalized medicine^1,2^. The NCI-MATCH trial in the United States reported an objective response rate (ORR) of 22% of ipatasertib in patients with AKT1 E17K mutations, and another 56% of the patients achieved disease stability^3^. In India, a study of 797 non-small cell lung cancer (NSCLC) patients carrying targeted driver gene mutations showed that receiving genotype-matched targeted therapy increased the median overall survival (mOS) from 9.3 months to 26.7 months^4^. Similarly, the Australian Molecular Screening and Therapeutic program found that patients received matched treatment after comprehensive genomic profiling (CGP) experienced longer survival (mOS 21.2 vs 12.8 months) than those received unmatched therapy^5^. Various clinical trials demonstrated significant clinical activity with precision oncology therapeutic approaches for several tumor types ^6^. Xu et al reported that in triple-negative breast cancer (TNBC), the median progression-free survival (PFS) was 5.7 months for the patients treated with TROP2 ADC Sacituzumab tirumotecan (SKB264) but was only 2.3 months for the patients treated with eribulin, vinorelbine, capecitabine, or gemcitabine (chemotherapy)^7^. Precision oncology showed optimistic responses in clinical trials at different stages. A study analyzed 224 articles related to a phase I oncology trial and found that the overall response rate (complete and partial responses) was 19.8%^8^. In a phase II clinical trial for patients with adenoid cystic carcinoma, the group treated with the targeted drug axitinib had a significantly longer median PFS (10.8 months) compared to the observation group (2.8 months)^9^. A placebo-controlled phase III trial showed that the risk of death in the sipuleucel-T group, Sipuleucel-T was used to treat metastatic castration-resistant prostate cancer, was relatively 22% lower than that in the placebo group^10^.

Precise drug efficacy evaluation is a fundamental prerequisite for precision medicine and anti-cancer drug screening^11^. While genomic profiling has become mainstream approach for matching targeted therapies in precision medicine, this approach exhibits substantial limitations. In the NCI-MATCH trial, the overall response rate among all evaluable patients is only 10.3%^12^. Tumor heterogeneity has been revealed as a main factor in the failure of anticancer therapies^13^. Therefore, personalized precise drug efficacy evaluation is becoming a necessary for individual patient ^11^. Three kinds of methods have showed great potential in facilitating drug selection: (1) Patient-derived tumor xenografts (PDXs) method allow an accurate prediction of tested drug efficacy, since they retain the idiosyncratic characteristics of different tumors from individual patients but are time-consuming (about 3 months from PDX model development to drug evaluation)^14,15^; (2) Patient-derived organoids (PDOs) method have been developed from a variety of cancers^16^. In the SENSOR and APOLLO studies, a failure rate (43% and 32%) was observed in organoid generation, which required an average of 10 weeks to develop, with drug screening taking an additional 8 weeks^17,18^; (3) Cell-based method typically involves the assays of the drug on cell viability, cell proliferation, colony formation, cytotoxicity, and so on, as well as the analysis cell morphological features^19^. Compared with PDXs and PDO, the cell-based method has advantages in terms of time and cost^20^.

Live cell high-content screening (HCS) is currently the main drug activity screening technology in oncology^21,22^. HCS utilizes microscopic imaging and quantitative image analysis technologies to obtain multi-dimensional cellular phenotypic features analysis^23,24^. Commercialized HCS microscope, such as Thermo Fisher Scientific’s CellInsight HCS platform, has been widely used in drugs efficacy evaluation and pharmaceutical industry^25–27^. Major advancements in HCS over the past five years include the application of deep learning, the emergence of label-free brightfield imaging, improvements in data analysis methods, the availability of large public datasets, and the integration of multi-modal data^28^. For example, a weakly supervised convolutional neural network (CNN) DML model was used to identify blood cancer cell morphology from bright-field and DAPI images, successfully screening 136 drug regimens and extending patients’ progression-free survival by 3.2 months^29^.

Based on the characteristic of fluorescence resonance energy transfer (FRET) microscopy in acquiring both cellular phenotypic information and the structural information of drug target proteins, we first develop a quantitative drug efficacy evaluation method (DL-TCP-FRET) which integrates drug target proteins interaction information into time- and concentration-dependent phenotypic features for the first time. We employ FRETscope to acquire both time- and concentration-dependent subcellular organelle phenotypic features, and FRET efficiency (*E*_*D*_) between target molecules of each living cells. Our method performed a dual-logistic (DL) analysis on the time- and concentration-dependent phenotypic features to establish a phenotypic efficacy score (*P* score). The acquired FRET efficiency (*E*_*D*_) between target molecules is used to construct a target molecule efficacy score (*T* score). Finally, *T* score was integrated into the *P* score to establish a comprehensive efficacy score (*PT* score). DL-TCP-FRET was performed for the NSCLC A549 cell line coexpressing CFP-EGFR and YFP-GRB2 (two targets) to evaluate the efficacy of six compounds including gefitinib (GEF), afatinib (AFA), dacomitinib (DAC), osimertinib (OSI), almonertinib (ALM) and vinorelbine (VIN). The experimental results demonstrated that DL-TCP-FRET not only distinguished targeted drugs and the non-targeted chemotherapeutic agent VIN but also obtained consistent efficacy of targeted drugs with the published research.

## 2. Materials and Methods

### 2.1 Reagents and antibodies

Geftinib (GEF, HY-50895A), Afatinib (AFA, HY-10261A), Almonertinib (ALM, HY-112823) and Osimertinib (OSI, HY-15772) were purchased from MedChemExpress (New Jersey, USA). Dacomitinib (DAC, A119719) and Vinorelbine (VIN, A676353) were purchased from Ambeed (Chicago, USA). Mitotracker Deep Red (M22426) was purchased from Thermo Fisher Scientific (Massachusetts, USA). Hoechst 33342 (C1022) was purchased from Beyotime (Shanghai, China). ExFect transfection reagent (T101-01) was purchased from Vazyme (Nanjing, China). Cell counting Kit-8 (GK10001) was purchased from Good Laboratory Practice Bioscience (California, USA).

### 2.2 Plasmid expression and vector construction

EGFR-CFP (G45807), GRB2 (P49764) and shRNA against EGFR (P69675) were purchased from Miaoling Biotech (Wuhan, China). GRB2-YFP plasmid was constructed by ourselves. Full-length GRB2 was released from the pCMV-GRB2(human)-3×HA-Neo plasmid by double digestion with XhoI and EcoRI, and inserted into the Bad-YFP plasmid using homologous recombination.

### 2.3 Cell culture and transfections

The A549 cells was obtained from Shangen (Wuhan, China). Cells were maintained in Roswell Park Memorial Institute 1640 medium (RPMI 1640, Boster, Wuhan, China), with 10% fetal bovine serum (FBS, BIOVISTECH, SE100-011), 100 U/ml penicillin G and 100 μg/ml streptomycin in a humidified atmosphere with 5% CO_2_ at 37 ℃. Cells were passaged early and cryopreserved and maintained in culture for <3 months, treated regularly for mycoplasma contamination using the luciferase mycoplasma detection kit (Transgen, Beijing, China). For transfection experiments, cells were plated in 20 mm glass dishes and cultured overnight in RPMI 1640 with 10 % FBS. When the cells achieved about 80 % confluence, plasmid transfection was performed using ExFect transfection reagent following the manufacturer’s instructions, with a transfection duration of 24 h.

### 2.4 Cell viability assay

The cell viability was measured by Cell Counting Kit 8 according to the manufacturer’s protocol. In brief, Cells were planted in 96-well plates and incubated in RPMI 1640 containing 10 % FBS at 37 ℃ for 24 h. Then, the cells were treated with different concentrations of Gef, Dac, Afa, Osi, and Vin for 24 h. Subsequently, 100ul of serum-free RPMI 1640 containing 10 % CCK8 solution was added to each well, and incubated at 37 ℃ for 1 h, The absorbance was measured at 450 nm using a micro-plate reader. Cell viability was normalized compared to the control group.

### 2.5 Hoechst 33342 and Mitotracker Deep Red staining

Cells (1 × 10^4^ cells/well) were seeded into a confocal dish. The cells were washed three times with 1×PBS and then stained with Hoechst 33342 and Mitotracker Deep Red (20 μg/ml at 37℃, 5% CO_2_) for 15 min under light-protected conditions, and then rinsed with 1×PBS three times prior to imaging.

### 2.6 Imaging system

A multi-modal quantitative FRET microscope (IX73, Olympus, Japan) was utilized as the imaging system for our method^30^. The microscope was equipped with a 60×/1.42NA oil lens (Olympus, Japan) objective and a CMOS camera (Flash 4.0, Hamamatsu Photonics, Japan); a bright-field LED light source; and the excitation light source was a 130W mercury lamp (U-HGLGPS, Olympus, Japan), including six intensity settings of 100%, 50%, 25%, 12%, 6%, and 3%. Five images were acquired from a specific field of view (FOV). The DD, DA, and AA images collectively constitute the E-FRET dataset of this FOV, with the remaining two being the nuclear and mitochondrial fluorescence image corresponding to the same FOV. For the details of imaging filter sets, see Table 1.

**Table 1.** Imaging filter sets.

| Channel | Excitation filter | Emission filter |
| --- | --- | --- |
| DD | 436/20nm | 480/20nm |
| DA | 436/20nm | 530/20nm |
| AA | 510/20nm | 530/20nm |
| Hoechst | 365/10nm | 440/40nm |
| Mitochondrion | 630/20nm | 660/20nm |

### 2.7 DL-TCP-FRET

As shown in Figure 1, DL-TCP-FRET consisted with four steps: (a) drug treatment and microscopic imaging; (b) FRET image processing and target molecule efficacy score (*T*) calculation; (c) cellular phenotypic image processing and phenotypic efficacy score (*P*) calculation; (d) comprehensive efficacy score (*PT*) calculation and quantitative efficacy evaluation of drugs. The details of these four steps are described as follows.

**Figure 1.**
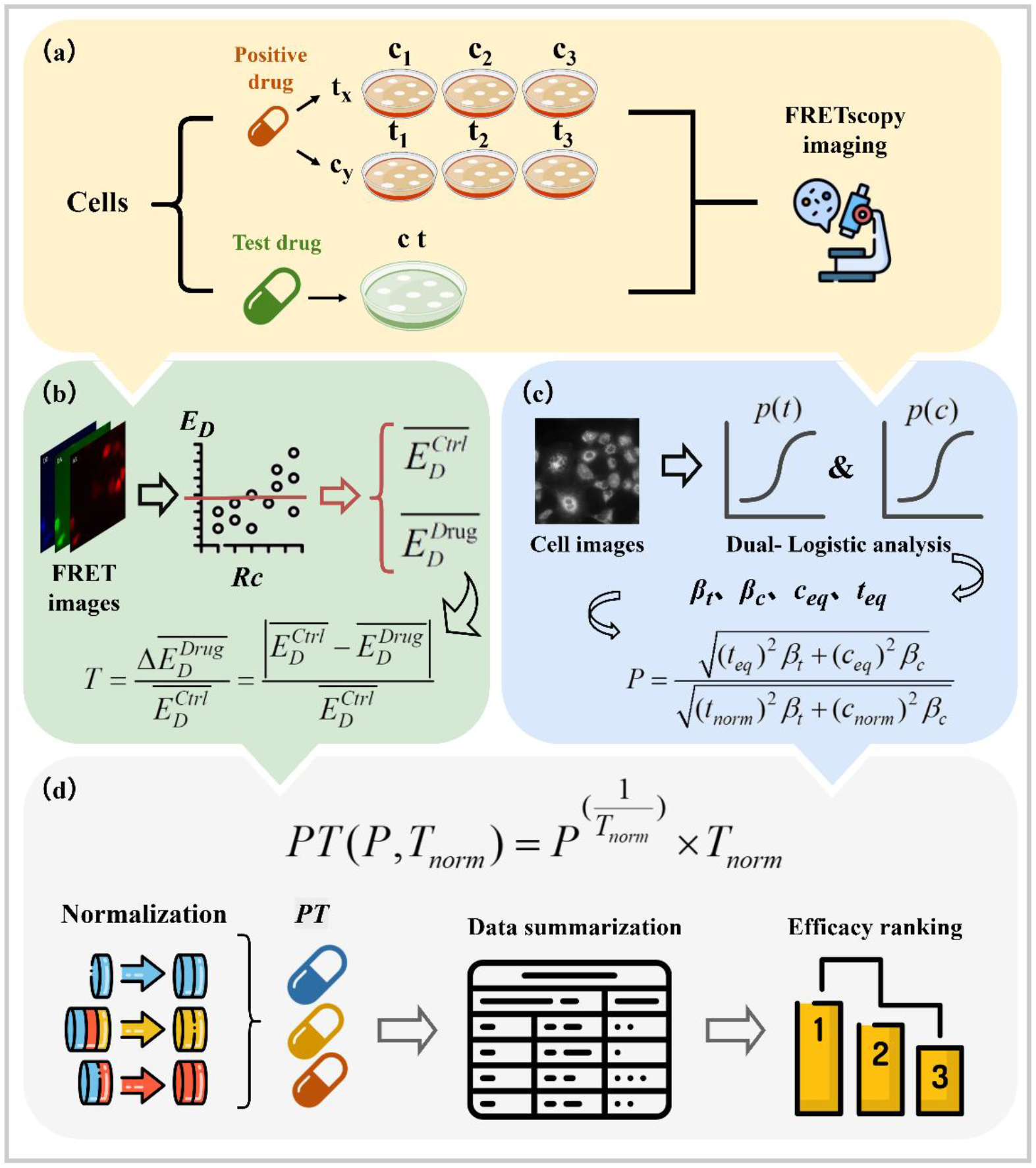
Flowchart of DL-TCP-FRET. (a) Drug treatment and microscopic imaging. (b) FRET image processing and T score calculation. (c) Cellular phenotypic image processing and P score calculation. (d) PT score calculation and quantitative efficacy evaluation of drugs.

#### 2.7.1 Drug treatment (Figure 1a)

Cells transfected with the target proteins were treated with drugs and divided into two groups. One group was administered a positive drug with set time gradients (fixed concentration) and concentration gradients (fixed time) to establish the dual-Logistic model. The other group was treated with six test drugs at any concentration and time. All samples are imaged by FRET microscope.

#### 2.7.2 FRET imaging-based T score (Figure 1b)

To acquire the target information of drug treatment, we selected the regions of interest (ROI) from the obtained FRET three-channel images (DD, DA, AA), and then calculate the donor-centric FRET efficiency (*E*_*D*_) and the donor-acceptor concentration ratio (*R*_*C*_) for each ROI, the corresponding *E*_*D*_ values were then selected based on a defined *R*_*C*_ range^31^. Subsequently, the mean *E*_*D*_ was calculated for both the drug treated group and the control group. Finally, the target efficacy *T* score is defined as:

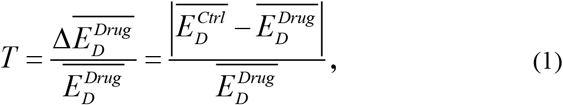

Where 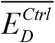 and 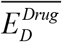 are the mean ED for control group and drug treated group, respectively.

Further, we performed a secondary normalization on the raw *T* scores to obtain a normalized *T* score (*T*_*norm*_). Considering that the FRET efficiency between target proteins is theoretically non-zero, the raw *T* score cannot reach the theoretical maximum of 1. Therefore, we used the maximum *T* score across all samples as the normalization reference to linearly map *T* scores to the 0-1 range, eliminating baseline variations across different batches or experimental conditions and enhancing the comparability of target scores.

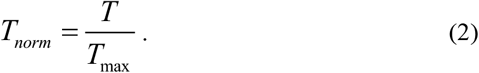

#### 2.7.3 Phenotypic feature-based P score (Figure 1c)

##### (1) FRET signal-guided Cell screening

Based on the acquired quantitative FRET three-channel fluorescence images (DD/DA/AA), we screened cells co-expressing both donor and acceptor proteins for each field of view. A separate mask was generated for each field of view, which was used to extract the phenotypic features of cells with protein co-expression (Figure 2).

**Figure 2.**
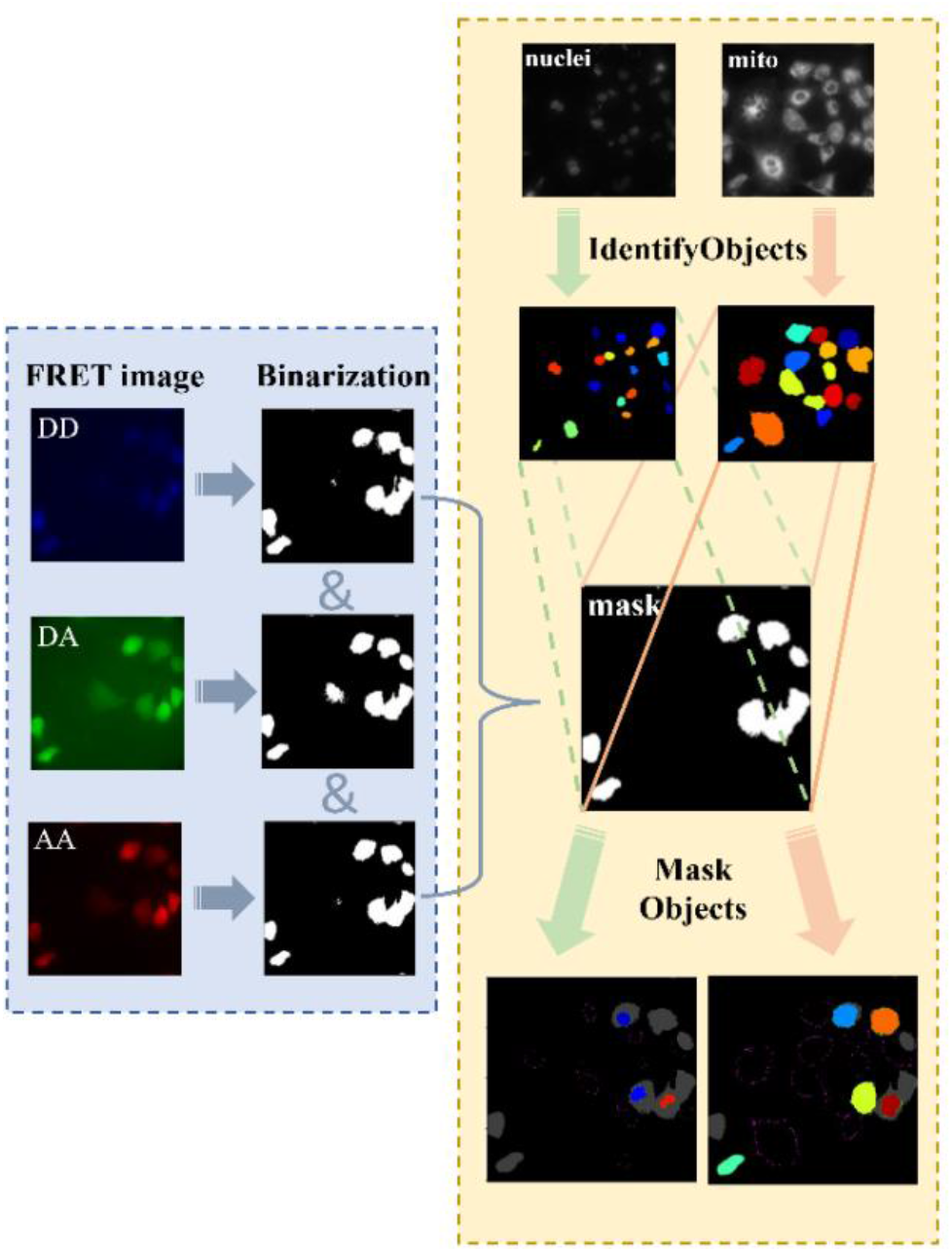
Flowchart for extracting phenotypic features of the cells co-expressing FPs-labelled donor and acceptor targeted proteins.

##### (2) Cellular phenotypic feature extraction

The cellular phenotypic features are extracted from nuclei and mitochondria fluorescence images using CellProfiler. The analysis pipelines are listed in Table 2. Using the CellProfiler software, nuclei and mitochondria fluorescence images from the same FOV were processed for noise removal and contrast enhancement. The processed nuclear and mitochondrial fluorescence images were then segmented and subjected to object recognition. Finally, the previously generated mask was applied to recognition results to screen out the nuclei and mitochondria from cells co-expressing both donor and acceptor proteins. Phenotypic features (e.g., shape and intensity) were extracted, and the data were output.

**Table 2.** CellProfiler pipeline and its functions for extracting cellular phenotypic features.

| Pipeline | Function |
| --- | --- |
| Threshold | Produces a binary, or black and white images. |
| ImageMath | Performs mathematical operations on images. |
| ReduceNoise | Reduces image noise and smooths images. |
| RescaleIntensity | Adjusts image pixel intensity range. |
| IdentifyPrimaryObjects | Identifies and segments primary objects. |
| MaskObjects | Uses a mask to retain or mask image regions. |
| MeasureObjectIntensity | Measures pixel intensity features of objects. |
| MeasureObjectSizeShape | Measures size and shape features of specified objects. |
| ExportToSpreadsheet | Exports all measurement results to a spreadsheet file. |

#### 2.7.4 Integration of phenotypic features

##### (1) Feature screening

For the modeling group treated with the positive drug under different time and concentration gradients, shape and intensity phenotypic features of the cells co-expressing FPs-labelled drug target molecules were extracted for each time and concentration using CellProfiler. First, we eliminated the phenotypic features that represent numbers, locations and directions. After outliers were eliminated through data cleaning (IQR), time-gradient curves and concentration-gradient curves corresponding to each of the features were obtained. We calculated the correlations between each feature and drug treatment time (*r*_*Ft,t*_) as well as between each feature and drug concentration(*r*_*Fc,c*_), respectively:

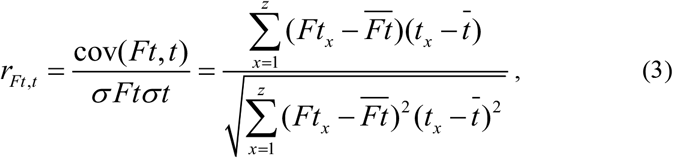

where *Ft*_*x*_ is the mean value of the *F* feature at drug treatment time *t*_*x*_, 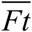 is the mean value of the *F* feature at all drug treatment time, 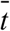 is the mean value of all the time-gradients, *z* is the number of all the time-gradients set up,

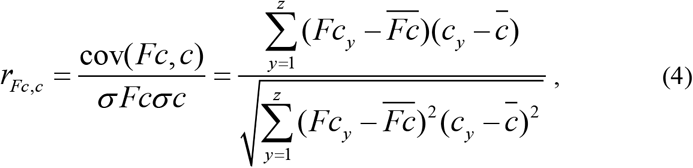

where *Fc*_*y*_ is the mean value of the *F* feature at drug concentration *c*_*y*_, 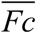 is the mean value of the *F* feature at all drug concentration, 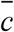 is the mean value of all the concentration -gradients, *z* is the number of all the concentration -gradients set up. Phenotypic features with |*r*_*Ft,t*_|+ |*r*_*Fc,c*_|≥1.8 and |*r*_*Ft,t*_|≥0.9 and |*r*_*Fc,c*_|≥0.9 were screened out as phenotypic features for drug efficacy evaluation in our method.

##### (2) Integration of phenotypic features (Figure 3)

For each phenotypic feature (such as *F* feature), the weight (*W*_*F*_) of the *F* phenotypic feature as follow:

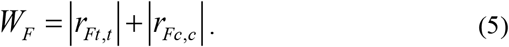

**Figure 3.**
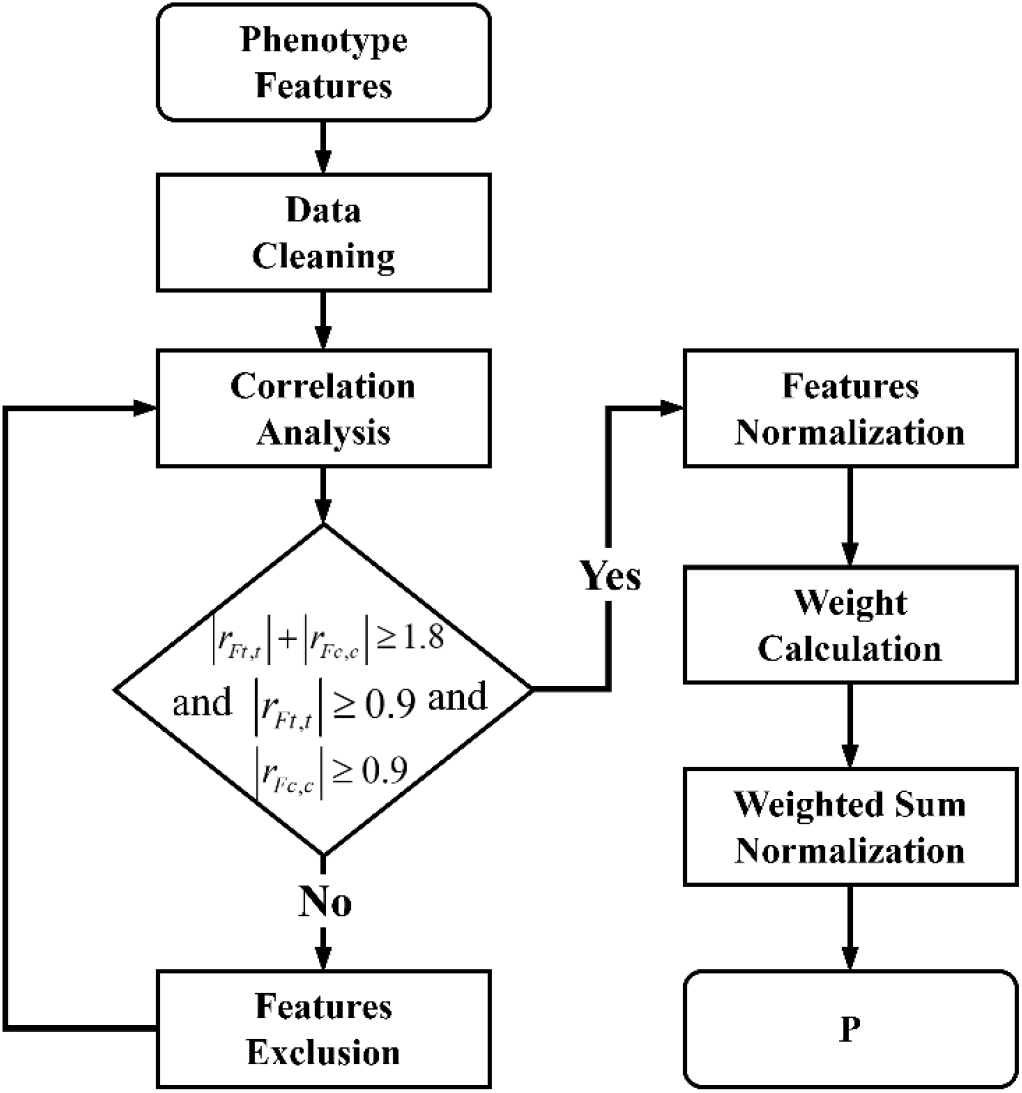
Flowchart of phenotypic characteristic value (p) calculation.

For each phenotypic feature (such as *F* feature), the maximum *F* value across modeling samples treated with positive drug is recorded as *F*_*max*_, the minimum *F* value across modeling samples treated with positive drug is recorded as *F*_*min*_, the normalized phenotypic feature value (*F*_*norm*_) of drug treated samples as follow:

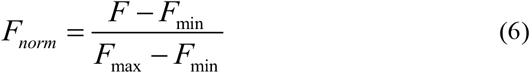

Finally, the phenotype characterization value *p* of each drug treated sample is:

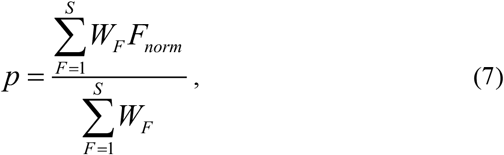

where *S* is the number of features were screened out.

For the modeling group treated with the positive drug, we can calculate the different *p* values corresponding to each drug treatment time and concentration gradient using Eq. (5). Similarly, we calculated six *p* values for the six test drugs using the same *S* features obtained through screening.

##### (3) Logistic fitting of time- and concentration-dependent phenotype

For a fixed drug treatment time, concentration-dependent *p* is used to perform Logistic fitting:

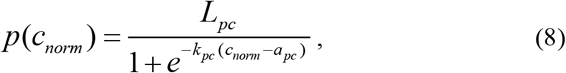

where 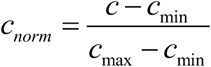 is the normalized concentration value of drug treated samples. Then the concentration-related phenotypic change growth rate parameter *k*_*pc*_, offset parameter *a*_*pc*_, and upper asymptote parameter *L*_*pc*_ were obtained.

For a fixed concentration, time-dependent *p* is used to perform Logistic fitting:

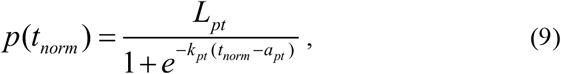

where 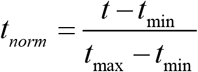 is the normalized drug treatment time value of drug treated samples. Then the time-related phenotypic change growth rate parameter *k*_*pt*_, offset parameter *a*_*pt*_, and upper asymptote parameter *L*_*pt*_ were obtained.

##### (4) P score Calculation

To unify the units, we perform normalization on the obtained parameters. 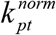 is defined as the ratio of *k*_*pt*_ to the maximum of *k*_*pt*_ and 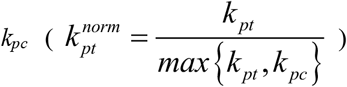. Similarly, normalized 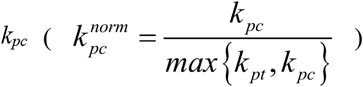, normalized 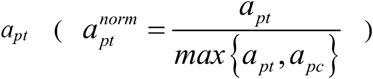, normalized 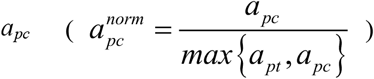, normalized 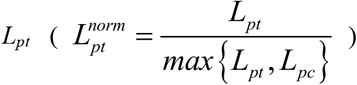 and normalized 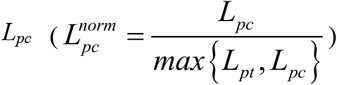 were calculated.

To balance the time- and concentration-dependent drug efficacy, we defined two vectors 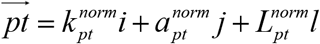 and 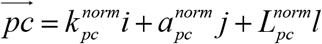. Therefore, the time weight *β*_*t*_ and concentration weight *β*_*c*_ are:

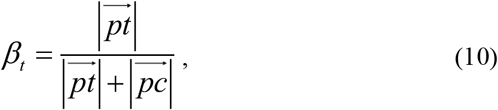

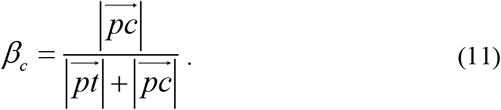

where 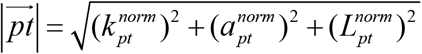 and 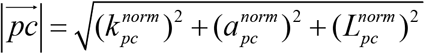 are the modulus of 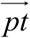 and 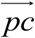, respectively.

Focusing on the time- and concentration- dependence of drug efficacy, this study is the first to adopt dual-logistic (DL) analysis for phenotypic feature modeling, calculating time weight (*β*_*t*_) and concentration weight (*β*_*c*_) to quantify their contributions to drug efficacy.

When the parameters kpt, apt, npt are fitted as constant, we can get the corresponding concentration (ceq) of a test drug from Eq. (7) with its phenotypic characterization value p. Similarly, we can get teq of a test drug from Eq. (8).

The fitted parameter values were substituted into the time-concentration dual Logistic model, and the value p of the test drugs was also substituted into the function. Next, performed the reverse mapping operation. Subsequently, solve for the equivalent normalized time (teq) and equivalent normalized concentration (ceq) of the test drugs.

The teq and ceq were weighted respectively to calculate the Euclidean distance, and the distance was then normalized to derive the phenotypic efficacy score (P):

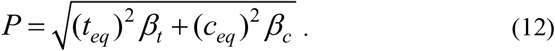

#### 2.7.5 PT score integrating the P and T scores

The *T* score and *P* score are integrated to obtain the comprehensive efficacy score *PT*.

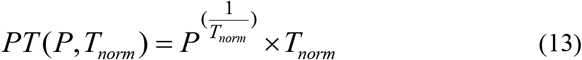

The *PT* score of all test drugs was normalized to make the results clearer.

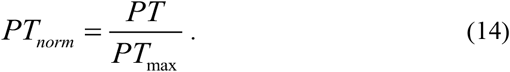

Similarly, normalized *P* score 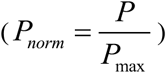 were calculated.

## 3. Results

Afatinib (AFA) was selected as the positive drug. For the concentration-dependent model, the treatment time was fixed at 8 h, with concentration gradients set as 0 μM, 12.92 μM, 25.84 μM, and 38.76 μM. For the time-dependent model, the concentration was fixed at the half-maximal inhibitory concentration (IC_50_, 25.84 μM), with different drug treatment times set as 0 h, 4 h, 8 h, and 12 h. These two sets of experiments were conducted to establish a concentration-dependent and time-dependent dual Logistic model.

Six test drug groups were treated at a unified concentration of 25.84 μM for 4 h, and a control group was set under the same treatment time for comparison.

### 3.1 FRET imaging-based E_D_ and T score

To validate the DL-TCP-FRET method for drug efficacy assessment, we established an evaluation system in A549 cells co-expressing CFP-EGFR and YFP-GRB2, followed by treatment with six drugs. Figure 4a shows the representative of protein distribution of AFA-treated group of 4 hours, for details of other drug-treated groups, refer to Supplementary Material S1. AFA’s specific inhibition of EGFR tyrosine kinase activity reduces the phosphorylation-dependent recruitment of GRB2 to EGFR, diminishing the interaction between the two proteins. In the *E*_*D*_ and *R*_*C*_ pseudo-color images, the abnormal high values observed in individual cells are due to apoptotic cells. During the process of cell apoptosis, cell shrinkage and breakdown of organelles lead to abnormal aggregation of fluorescent proteins, thereby causing non-specific enhancement of *E*_*D*_. The apoptotic cells were excluded from the subsequent quantitative analysis to avoid interfering with the accuracy of the efficacy assessment.

**Figure 4.**
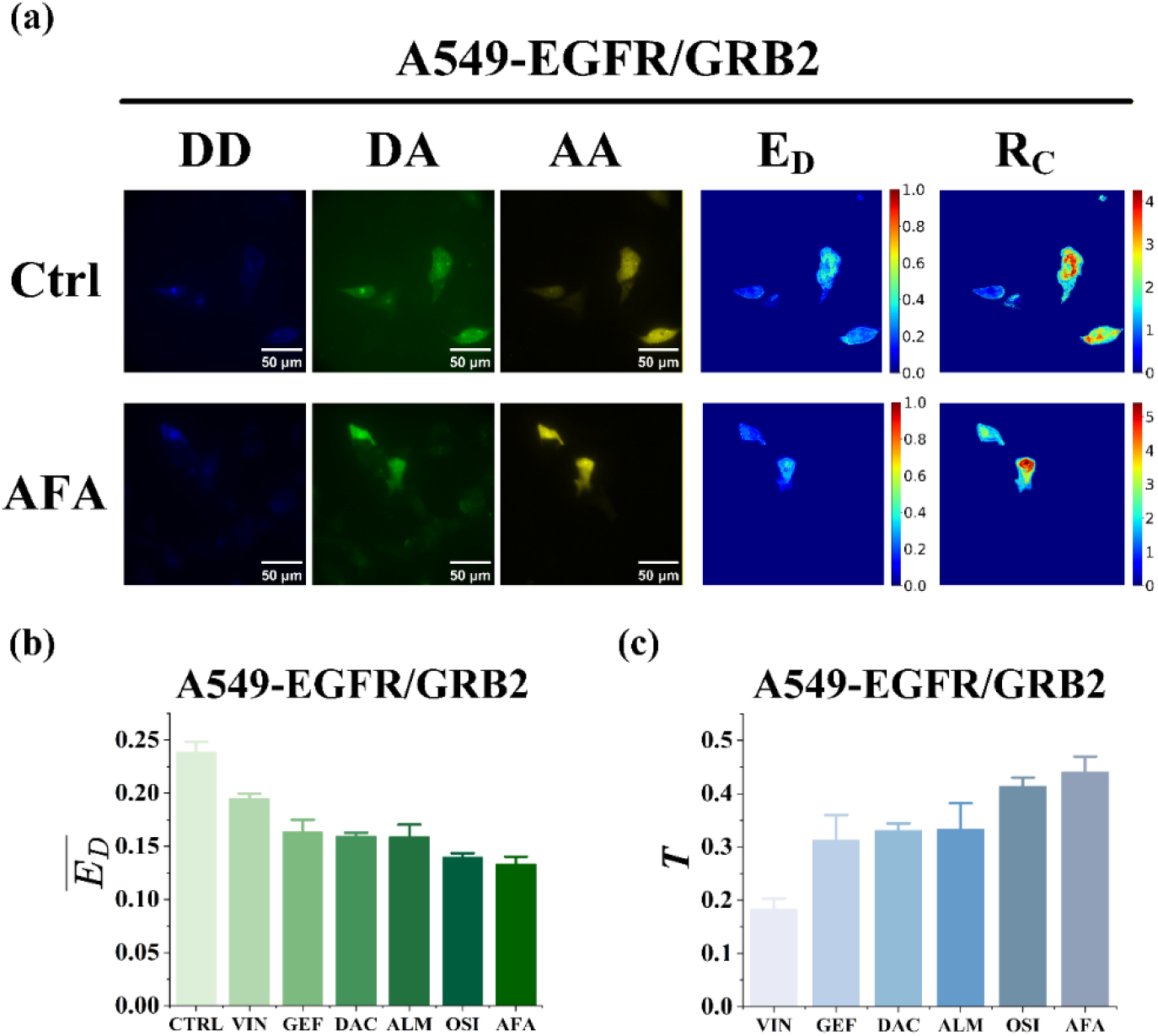
(a) Representative FRET three-channel images and *E*_*D*_, *R*_*C*_ pseudo color images of the AFA-treated group and the control group. (b) 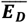 of six drugs. (c) *T* score of six drugs.

Quantitative analysis of FRET efficiency showed significant differences in *E*_*D*_ values between all drug-treated groups and the Ctrl group (Figure 4b). Specifically, the 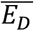 value of the AFA group was 0.13, significantly lower than the 0.27 of the Ctrl group, indicating that AFA dissociate the binding of GRB2 to EGFR. According to Eq. (1), the *T* score of AFA was |0.24-0.13|/0.25=0.44. Similarly, the *T* scores of ALM, OSI, DAC, GEF and VIN are 0.43, 0.41, 0.33, 0.31 and 0.18 (Figure 4c). Among six drugs, AFA had the highest T scores, indicating blocking effects on EGFR-GRB2 binding. In contrast, the *T* score for the chemotherapeutic agent VIN was significantly lower, as VIN is not a drug specifically targeting EGFR-GRB2.

### 3.2 Phenotypic feature-based P score

#### 3.2.1 Phenotypic features screening

206 phenotypic features were extracted from cell nuclei and mitochondria fluorescence images using CellProfiler. Then, we eliminated the phenotypic features that represent numbers, locations and directions. We screened out phenotypic features with |*r*_*Ft,t*_|+ |*r*_*Fc,c*_|≥1.8 and |*r*_*Ft,t*_|≥0.9 and |*r*_*Fc,c*_|≥0.9 after AFA treatment as phenotypic features for drug efficacy evaluation. These features are stable because they change in a single direction over time and with concentration. The five features which were screened out are shown in Table 3.

**Table 3.** Five phenotypic features and their rFt,t, rFc,c and weights in A549 Cell Line.

| Features | $r_{Ft,t}$ | $r_{Fc,c}$ | Weight |
| --- | --- | --- | --- |
| NU_AreaShape_Eccentricity | 0.90 | 0.98 | 1.88 |
| NU_AreaShape_Zernike_2_2 | 0.99 | 0.98 | 1.97 |
| NU_AreaShape_Zernike_4_4 | 1.00 | 0.98 | 1.98 |
| MITO_AreaShape_Extent | -0.98 | 0.98 | -1.96 |
| MITO_AreaShape_Zernike_9_7 | 0.90 | 1.00 | 1.90 |

**Table 4.** Five phenotypic features after AFA treatment at 25.84μM for 4 hours.

| Features | F | F<sub>max</sub> | F<sub>min</sub> | F<sub>norm</sub> |
| --- | --- | --- | --- | --- |
| NU_AreaShape_Eccentricity | 0.0123 | 0.0743 | 0 | 0.17 |
| NU_AreaShape_Zernike_2_2 | 0.0020 | 0.0076 | 0 | 0.26 |
| NU_AreaShape_Zernike_4_4 | 0.0014 | 0.0037 | 0 | 0.37 |
| MITO_AreaShape_Extent | 0.0178 | 0.0745 | 0 | 0.24 |
| MITO_AreaShape_Zernike_9_7 | 0.0001 | 0.0018 | 0 | 0.06 |

#### 3.2.2 Phenotypic features integrating and p value calculation

The weights are calculated according to Eq. (4), and the weights of the five features are shown in Table 3. The representative images of the cell mitochondria and nuclei treated with AFA at the 25.84 μM for 4 hours are shown in Figure 5a. The *p* value of this group is 0.22, according to Eq. (6). Similarly, we can calculate seven *p* values in total, consisting of different time (0h, 4h, 8h, 12h) and concentrations (0 μM, 12.92 μM, 25.84 μM, and 38.76 μM). Features and *p* values for the other groups can be found in Supplementary Material S2.

**Figure 5.**
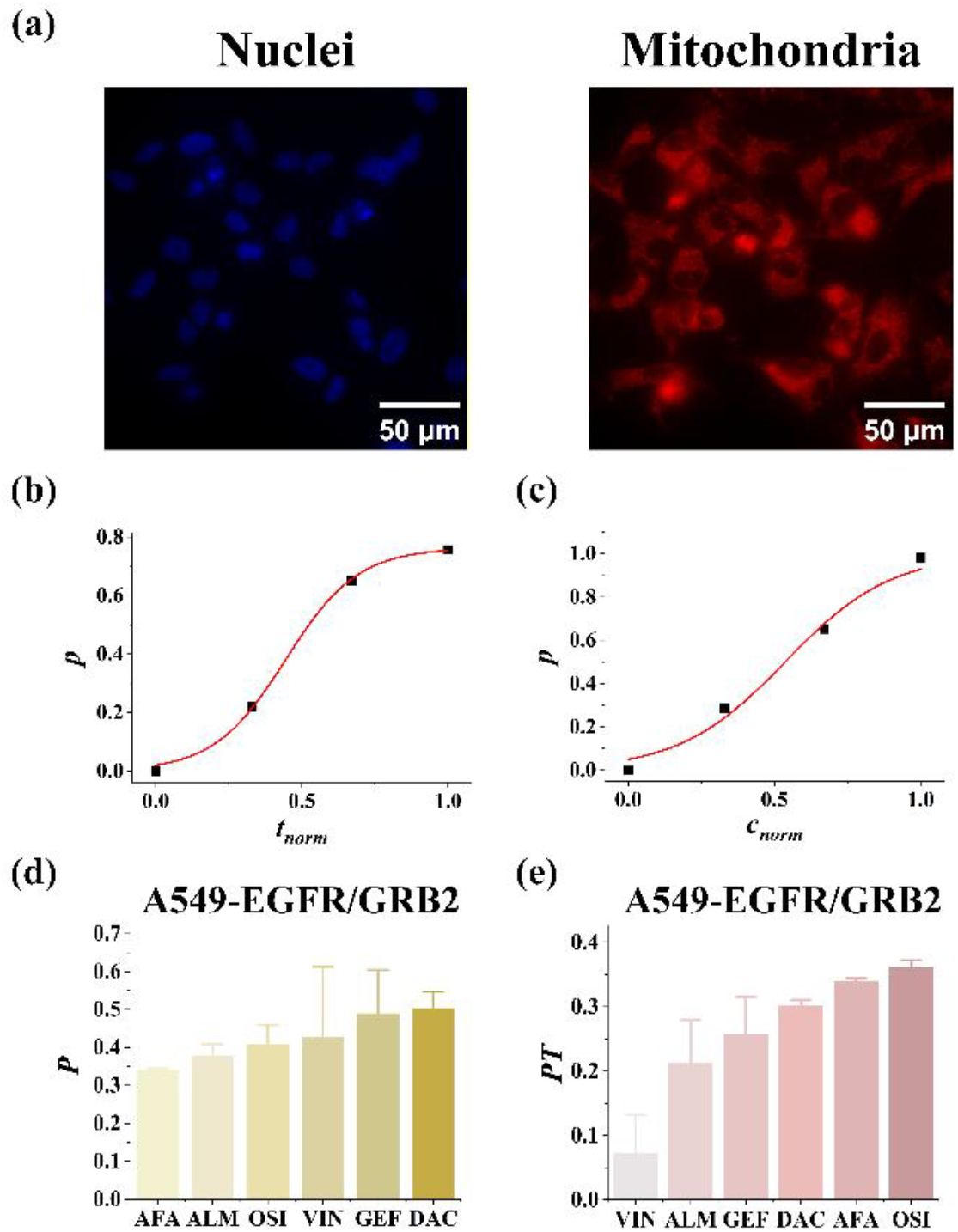
(a) Representative images of mitochondria and cell nuclei treated with AFA at 25.84 μM for 4 hours. (b) Logistic fit curve of *P* versus normalized time. (c) Logistic fit curve of *P* versus normalized concentration. (d) *P* score of six drugs. (e) *PT*_*norm*_ of six drugs.

Time-dependent phenotypic function and concentration-dependent phenotypic function were fitted respectively. The specific forms of the two functions are listed as follows:

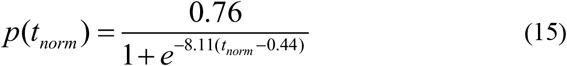

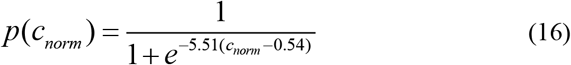

Parameters derived from the fitting are shown in Table 5. The fitting results are shown in Figure 5b and 5c.

**Table 5.** Parameters of phenotype changes in time-concentration-dependent models.

| Parameters | $k_{pt}$ | $a_{pt}$ | $L_{pt}$ | $k_{pc}$ | $a_{pc}$ | $L_{pc}$ |
| --- | --- | --- | --- | --- | --- | --- |
| Value | 8.11 | 0.44 | 0.76 | 5.51 | 0.54 | 1 |

The results revealed distinct response patterns between the two dimensions. The time dimension exhibited a faster phenotypic change rate than the concentration dimension (*k*_*Pt*_=8.11 vs *k*_*Pc*_=5.51). The offset parameter indicated that the normalized time required to trigger significant phenotypic changes was 0.44 (*a*_*Pt*_=0.44), and the normalized concentration corresponding to such changes was 0.54 (*a*_*Pc*_=0.54).

Phenotypic time weight (*β*_*t*_ = 0.49) and concentration weight (*β*_*c*_ = 0.51) were calculated using Eq. (9) and Eq. (10), with the *β*_*t*_ being approximately twice the *β*_*c*_ (Table 6). The *β*_*c*_ and *β*_*t*_ reflect the respective contributions of drug action time and drug concentration to phenotypic changes. During the process of EGFR-TKIs acting on A549 cells co-expressing CFP-EGFR and YFP-GRB2, the regulatory efficiency of concentration on phenotypic feature changes is significantly superior to that of drug treatment time.

**Table 6.** Parameters of time weight and concentration weight.

| Parameters | $\beta_t$ | $\beta_c$ |
| --- | --- | --- |
| Value | 0.49 | 0.51 |

**Table 6.**
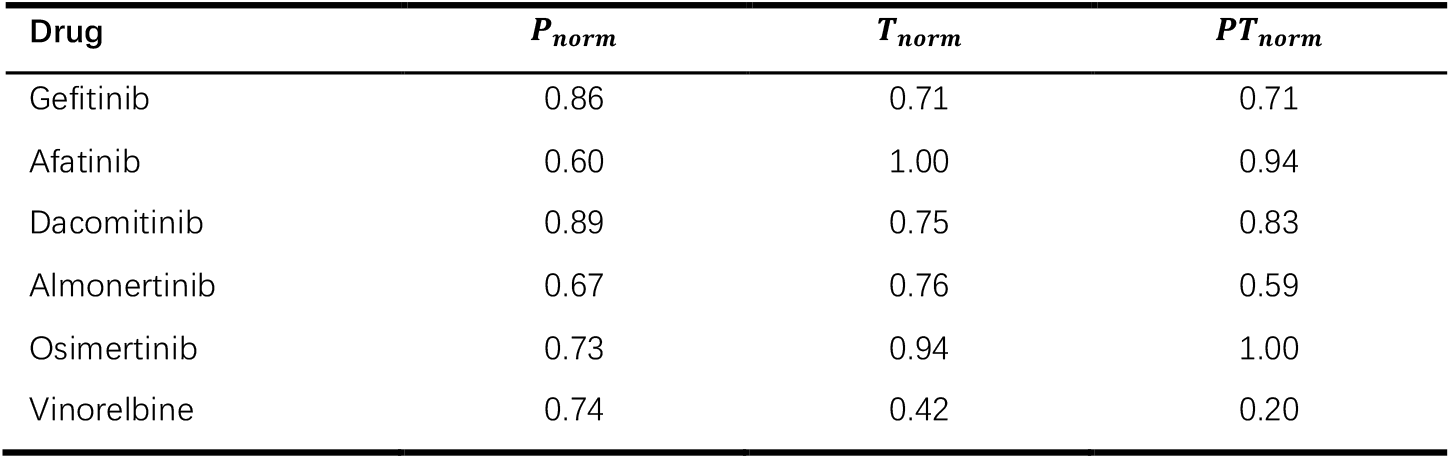
Normalized *P, T* and *PT* scores of six drugs.

The value *p* of the six drugs was substituted into Eq. (14) and Eq. (15), followed by a reverse mapping operation. The equivalent normalized time (*t*_*eq*_) and equivalent normalized concentration (*c*_*eq*_) of the test drugs were solved. Finally, the *P* scores of the six drugs were calculated from Eq. (11) to quantify efficacy in regulating EGFR-related phenotypes (Figure 5d). For example, the *c*_*eq*_ of AFA-treated group is 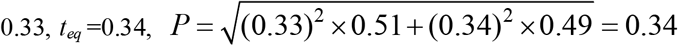. For details of other drug-treated groups, refer to Supplementary Material S3. The *P* score values for other test drugs are shown in Figure 5d.

#### 3.3 P- and T-based PT score of EGFR-TKIs and chemotherapeutic agent

*PT* scores were obtained by integrating P scores and T scores using Eq. (12) (Figure 5e). A significant distinction in *PT* scores was observed between chemotherapeutic agents and other EGFR-targeted drugs. VIN is a microtubule inhibitor and chemotherapeutic agent which mainly exerts broad-spectrum cytotoxicity by blocking mitosis in tumor cells and does not have the targeting specificity for the EGFR target and signaling pathway. For the chemotherapeutic agent VIN, the *P* score was 0.20 and the *T* score was 0.18. However, the *PT* score of VIN calculated by the established model approached 0. The results demonstrate our model can distinguish non-specifically targeted drugs. To compare the drug efficacy more clearly, we normalized the *P, T* and *PT* score in Table 6. The OSI ranked highest, followed by AFA, DAC, GEF, ALM, and finally VIN.

## 4. Discussion

The DL-TCP-FRET integrates drug target information with time- and concentration-dependent phenotypic features to quantitatively evaluate the efficacy of anticancer drugs. Since drug efficacy is a time- and concentration-dependent variable32, we selected the Logistic model as the foundation of our method, as it can appropriately describe the density-dependent growth pattern of cells characterized by initial exponential growth, intermediate maximum growth rate, and late saturation33. Furthermore, this pattern is consistent with the S-shaped dose–response curve (EC_50_) commonly used in pharmacodynamics and Pasetto et al. mathematically demonstrated that the Logistic model is superior to the simple exponential model in tumor cell growth mechanism consistency, parameter identifiability and early predictive performance34.

Based on the Logistic model, we established a dual-logistic (DL) model incorporating both time-dependent and concentration-dependent effects for the NSCLC A549 cell line with the EGFR/GRB2 target. Afatinib, an EGFR-TKI with well-defined efficacy, was used as the reference drug, and a universal evaluation model was constructed using only one complete set of time-concentration gradient experiments on this reference. For other test drugs, no full gradient experiments were required, and rapid efficacy quantification could be achieved with only a small number of experimental data points. The DL-TCP-FRET simplifies the experimental workflow, reduces drug and cell consumption, improves the stability and comparability of results, and enhances the efficiency of in vitro anticancer drug screening.

Though the target score (*T*) and phenotypic score (*P*) are calculated independently, to screen drugs with target specificity, namely effective drugs that triggers both target response and phenotypic change simultaneously (an AND relationship), we initially used *PT(P,T)=P×T*, then optimized it to *PT(P,T)=P*^*(1/T)*^*×T* to avoid false positives caused by low *T* but excessively high *P*. The term *P*^*(1/T)*^ dynamically modulates *P* score according to *T* score: the higher *T* is, the smaller 1/T is, and the larger *P*^*(1/T)*^ *is*, thereby enhancing the phenotypic contribution with increasing target strength. This improvement effectively reduces false-positive results derived from non-target-dependent effects and ensures that only drugs with both significant target response and phenotypic inhibition can achieve high *PT* scores, making our method more accurate in identifying anticancer drugs with genuine target specificity. Notably, the *T* score was replaced by *T*_*norm*_, which calibrates the FRET baseline and ensures normalization consistency.

In the A549 cell line, the DL-TCP-FRET can reliably distinguish the non-targeted chemotherapeutic agent VIN (not an EGFR specific inhibitor^35^) from other EGFR-TKIs (AFA, OSI, DAC, ALM and GEF). We obtained targeted drugs’ efficacy results roughly consistent with the published research. A TMT-10plex phosphoproteomic study validated AFA and OSI as EGFR pathway inhibitors^36^, and a colony-forming assay showed that OSI exhibited stronger anti-tumor activity than AFA in A549 cells. DAC, ALM and GEF showed higher efficacy than VIN. These results verify the reliability of our method for evaluating drug efficacy at the cellular level.

Similar to Kornauth et al.’s EXALT study^37^, our method aims to quantify ex vivo drug efficacy using patient or cell samples, but differs from their scFPM protocol. While scFPM uses single-cell survival or death as the primary drug efficacy endpoint, the DL-TCP-FRET integrates subcellular phenotypes and protein-protein interactions, achieving the unification of “mechanism-driven” (target verification) and “result-driven” (phenotypic changes).

Despite its advantages in quantitative drug efficacy evaluation, the DL-TCP-FRET still has certain limitations. Firstly, the validation was directed at the adherent NSCLC A549 cell line, while clinical samples exhibit greater heterogeneity and complexity. Future studies will validate the model in PDOs or primary tumor cells. Secondly, only one target pair (e.g., EGFR-GRB2) was studied, while tumorigenesis often involves dynamic compensation and coordinated action of multiple signaling pathways, subsequent studies will develop a model for synchronously evaluating the interactions of multiple targets within single cells. Thirdly, only single-drug efficacy was evaluated. Thus, the DL model needs to be optimized to quantify the synergistic effect of combined drugs. Furthermore, integrating automated real-time imaging systems with deep learning-based morphological analysis will make the DL-TCP-FRET more accurate and reliable.

## 5 Conclusion

In conclusion, the DL-TCP-FRET integrates drug target information with time-dependent and concentration-dependent phenotypic characteristics for the quantitative evaluation of anticancer efficacy. This method not only simplifies the experimental workflow by reducing redundant time and concentration gradient experiments, but also effectively eliminates false-positive results through dynamic modulation of phenotypic scores based on target signals, enabling clear distinction between EGFR-TKIs and non-targeted chemotherapeutic agents. Our findings are consistent with published studies, verifying the reliability and feasibility of the method at the cellular level. Although certain limitations remain to be addressed, the DL-TCP-FRET method provides a new technical tool for preclinical drug screening with high efficiency and specificity. It is expected to promote the development of precision medicine and accelerate the discovery of new targeted anticancer drugs by bridging the gap between drug-target interactions and cellular phenotypic responses.

## Authorship contribution statement

Lingli Wang: Writing – original draft, Visualization, Software, Experiment, Methodology, Data curation, Conceptualization. Rumeng Qu: Writing – original draft, Experiment, Supervision. Qialing Huang: Writing – Experiment, Supervision. Min Hu: Writing – review & editing, Validation, Supervision, Conceptualization. Tongsheng Chen: Writing – review & editing, Supervision, Methodology, Conceptualization.

